# Perceptual learning mechanisms with single-stimulus exposure

**DOI:** 10.1101/2025.06.25.661650

**Authors:** María del Carmen Sanjuán, James Byron Nelson

## Abstract

The present manuscript briefly reviews associative and non-associative theories of perceptual learning and presents two experiments (1a & 1b) examining the extent to which retrospective revaluation (RR) and a restoration in perceptual effectiveness (RPE) can account for the enhanced discriminability that experience with a single stimulus produces. In both experiments participants were exposed to compound visual stimuli BX, CY, and DZ in an online video-game method followed by conditioning with compound stimuli AX and AY. In Experiment 1a generalization to compounds BX, DX, BY, and DY was assessed. Generalization to compounds involving B, either BX or BY, was less than to DX or DY, possibly reflecting enhanced salience of B, or that B had become inhibitory for the outcome through RR. Experiment 1b used a retardation test comparing compound stimuli BW and DW. Acquisition was more effective with BW than DW, suggesting that B had become more perceptually effective, resulting in it producing more external inhibition than D in Experiment 1a, and allowing it to condition more rapidly in Experiment 1b. Results are discussed with respect to current theories of perceptual learning and are consistent with that offered by Hall [17].

## 1 Introduction

### 1.1 Perceptual learning mechanisms in single-stimulus exposure

In our daily lives, we learn to distinguish between subtly different stimuli, even when we have direct experience with only one of them. For example, we can recognize a new coffee as different from other we have tasted before. If a particular food produces a gastric illness that leads to rejection of that food in the future, the conditioned aversion could lead to a rejection of a novel but similar-tasting food. Yet, if the similar food had been experienced prior to the illness episode, less or non-aversion can occur.

Experiencing a stimulus establishes a representation of that stimulus that allows it to be differentiated from other stimuli later. This pattern reflects perceptual learning: the processes involved in how the brain refines its capacity to identify and distinguish between similar stimuli based on even limited exposure. This article explores how exposure to a single stimulus can reduce generalization to similar stimuli, investigating the roles of retrospective revaluation (e.g. [11]) and recovery of perceptual effectiveness [17].

Perceptual learning studies are based on ideas that follow stimulus-sampling theory (e.g., [12],[13]) by assuming that stimuli are composed of many different elements, and generalization results from elements that stimuli have in common. They have predominately focused on the beneficial effects of exposing two stimuli made similar by the addition of a common explicit element, (e.g., AX and BX). When one stimulus is conditioned (AX+), substantial generalization to the test stimulus (BX) is observed. That generalization is assumed to largely result from the conditioning accrued to the common X elements. Prior exposure to these stimuli in alternation (e.g., AX/BX/AX/BX..) or in blocks (e.g., AX, AX,… BX, BX…) generally enhances the discrimination between AX and BX, with the intermixed type of exposure being more effective (e.g. [2]; [5],[6],[24],[64],[65],[72],[73]).

As with exposure to both the conditioned (AX) and test (BX) stimuli, exposure to only the test stimulus can reduce generalization ([3],[69],[61],[62],[63]). For instance, Sanjuan et al. [66] demonstrated that exposure to a flavor compound “BX” consisting of B (e.g., a solution of water with 1% salt) and X (e.g., hydrochloric acid) significantly reduced the generalization of an aversion conditioned to a similar flavor, AX, (where A is, for example, 10% sugar). The reduction with exposure to BX was more pronounced than with exposure to B and X separately. This effect, observed in both animals (e.g., [4],[72], [65],[66]) and humans [67], also represents a form of perceptual learning, yet it poses a challenge to both associative (e.g., [17],[38],[39]) and non-associative (e.g., [15]) theories. These theories, otherwise successfully explain perceptual learning effects that follow alternated exposure to both stimuli (AX and BX), and its superiority to blocked exposure.

### 1.2 Gibson’s (1969) non-associative account of perceptual learning

According to Gibson [15] perception improves with experience as individuals actively learn to differentiate and interpret sensory information. Such perceptual learning occurs through a process of differentiation, wherein the response to a stimulus becomes more specific and attuned to present stimulation with repeated exposure. As individuals encounter a stimulus more frequently, they refine their ability to detect invariances in the stimuli - those features that distinguish one stimulus from another-while ignoring irrelevant and non-informative features that do not help differentiate it from others.

In this process, the organism progressively refines its ability to detect the distinctive features of a stimulus while disregarding the shared elements that hinder differentiation, ultimately improving the stimulus’ discriminability from others. This differentiation is supported by attentional mechanisms, which are influenced by conditions that facilitate perceptual learning. Gibson, however, did not explicitly detail the precise mechanisms involved in differentiation; instead, her theory emphasized the importance of conditions—such as the opportunity for direct comparison—that promote their operation.

Gibson’s theory offers a ready explanation for the consistently observed intermixed-blocked effect in perceptual learning. The intermixed condition provides more opportunities for comparison between different stimuli, thereby enhancing discrimination and perceptual differentiation. In contrast, when stimuli are presented in a blocked sequence, the learning that occurs benefits less from the comparison mechanism, limiting the potential for perceptual improvement. Exposure to only the test stimulus BX, prior to conditioning of AX, allows for little opportunity for Gibson’s differentiation to operate. Nevertheless, repeated exposure to BX alone is sufficient to enhance its discriminability from other similar stimuli ([4],[3],[8],[25], [65],[66],[67], [73]).

### 1.3 Perceptual learning as inter-element associations

McLaren et al. ([38],[39]) proposed an associative mechanism for perceptual learning, based on the source of the activation of a stimulus representation. For instance, when AX is present, the representations of A and X are activated externally by the presence of the physical stimuli. However, the representation of the elements may also be “internally” activated by way of associations. When complex stimuli (e.g., AX and BX) are presented, it is assumed that bidirectional associations form between the individual elements X and A, as well as between X and B, a process referred to as “unitization.” These elements can also be associated with the background contextual stimuli present during their presentations. The result of these associations is that when one element is absent, the present element can retrieve or activate the internal representation of the other element. This unitization allows all elements of a complex stimulus to be internally active and present, even in when the entire stimulus is only partially present in the external environment.

Unitization occurs as the stimuli are repeatedly presented together. However, salience is assumed to be different for internally and externally activated stimuli, with internally activated stimuli being less salient. Internal activation is also assumed to limit the effectiveness of external activation therefore unitization leads to a reduction in the salience of individual elements X, A, and B, when presented in compounds and, consequently, diminishes the overall salience of the compound stimuli AX and BX. Following the establishment of these associations, McLaren and colleagues identify two primary mechanisms that contribute to perceptual learning; differential losses of stimulus effectiveness and mutual inhibition between unique elements.

### 1.4 Differential losses of stimulus effectiveness

The first mechanism involves a differential loss of effectiveness to the common elements (X) shared by two stimuli, AX and BX, compared to the unique A and B elements specific to each stimulus. The reduction in salience the model specifies to occur with internally activated stimuli is assumed to be a mechanism producing “latent inhibition,” where exposure to a simple CS subsequently retards conditioning with that stimulus (e.g., [35],[34],[46],[50], [20]). When AX and BX are pre-exposed, the X elements receive twice the exposure that A and B receive. That extra exposure results in stronger associations with the context, producing a stronger internally activated representation, and a greater reduction in the salience of X. During conditioning of AX, the A element, whose salience has been reduced to a lesser degree, acquires greater associative strength than X. Consequently, when BX is presented in a generalization test, the generalized response controlled by X is reduced compared to any condition where this differential loss of salience did not occur.

With respect to exposure to only the test stimulus BX, the differential latent inhibition mechanism cannot easily explain the results obtained by Sanjuan et al. [65]. In that study, one group of rats received four exposures to flavors AX and BX in alternation, another group received four exposures to AX and BX in separate blocks, and a third group received eight exposures to BX. Following aversive conditioning with AX, the consumption of BX was tested. Exposure to BX alone led to a greater reduction in generalization than exposure to both stimuli, whether presented in alternation or in separate blocks, despite that the amount of exposure to the X element was matched across all groups. That is, in terms of a reduction in salience of X due to its internal activation, the groups should have been equivalent.

Similar studies by Sanjuan et al. [66] also showed that exposure to BX was superior to conditions where the elements (B and X) had been separately exposed. In a second experiment in Sanjuan et al., [66] four different groups of rats varied with respect to the extent to which BX was exposed while matching exposure to X. Group 1 received 1 exposure to BX and 7 to X; Group 4 received 4 exposures to BX and 4 to X; Group 8 received 8 exposures to BX, while Group 0 received no exposure (i.e., exposure to water). If latent inhibition was the mechanism responsible for the BX exposure effect, no differences would be expected to occur. Generalization was not equivalent across the groups. Generalization was a function of the amount of exposure to the compound BX. The groups showed graded generalization where 1 exposure showed the greatest generalization, 4 some less and 8 the least. These results suggested that exposure to BX caused some type of perceptual learning about the stimulus that went beyond the responses governed by X.

The course of conditioning of AX was also revealing. All the groups receiving pre-exposure to BX received the same overall exposure to X, varying only by whether that exposure was with B. One or 4 exposures to BX (7 or 4 exposures to X alone) produced a delay in conditioning with AX. That is, any latent inhibition accrued to X generalized to AX during its conditioning phase. Yet, when all exposures to X were with B, there was no generalization to AX. Latent inhibition that accrued to X during its exposure within BX did not generalize well outside of the compound.

### 1.5 Mutual inhibition between unique elements

As discussed previously, a common finding in the literature is that alternated exposure is more effective than blocked exposure. The idea that associations form between the elements presented during pre-exposure gives rise to a second mechanism highlighted by McLaren et al. ([38],[39]) which accounts for the intermixed/blocked difference. When AX is repeatedly presented in alternation with BX, each presentation of AX may evoke the representation of B through its association with X. As B is thus “predicted” to be present, but physically absent, inhibition could form between A and B. During BX trials, A would similarly be evoked, facilitating an inhibitory connection between B and A. This latter connection is particularly relevant for reducing generalization, as it would operate during the BX test. On test with BX, inhibition between B and A would diminish any contribution that might arise from the associatively evoked image of the previously conditioned A by way of its association with X. The inhibitory connection between the present B and A would impede such a mediated effect from occurring.

### 1.6 Configural hypothesis

In the field of perception, it is widely discussed that exposure enables a highly precise, complete, and detailed encoding of a stimulus ([15],[19],[71],[16]). If the way a stimulus is encoded changes with experience, extended experience might produce a configural type of perception that makes the stimulus more dissimilar to other similar stimuli and its own components. Learning theories with some type of configural construct, such as those proposed by Pearce ([56],[57]) or Wagner [77]argue that when stimuli are presented together, the organism learns about the unique configuration of the compound rather than only learning about the individual components separately. A BX compound would be treated as a distinct stimulus configuration itself (Pearce) or contain a unique cue produced by the combination of the elements (Wagner), which might involve specific associative links between B and X that contribute to the overall configuration [39]. These perspectives emphasize that learning is not just about the simple addition of individual stimulus properties but about the interaction and relational binding of these properties.

This configural hypothesis may be relevant to understanding the effects of exposure to BX. If prolonged exposure causes BX to be perceived as a distinct stimulus, different from the sum of its components, generalization would be reduced. After exposure to BX, strong within-compound associations between B and X make BX distinct from other stimuli, such as AX, by emphasizing the unique combination of B and X as a single unit. BX would be perceived as a unique entity, different from its elements and different from those that include its elements like AX. This refined representation would mean that responses conditioned to AX do not readily transfer to BX.

That mechanism could explain the finding of Sanjuan et al., [66], discussed earlier, where 8 exposures to BX led to less latent inhibition in the conditioning of AX. Rodriguez and Alonso ([61] found direct evidence of exposure to BX reducing generalization by conditioning X directly, where generalization is not affected by the addition of an A element. After 8 exposures to BX, X conditioned better (i.e., showed less latent inhibition that may have accrued during its exposure with B) than after 4 exposures. Despite more exposure to X in the 8-exposure group, less latent inhibition was observed, suggesting that the increased exposure to BX led to the compound being perceived as a whole configural unit such that there was a failure to recognize X alone as having been experienced before.

Despite that findings showing preexposure to BX reducing generalization to X are consistent with the idea that BX exposure leads to it being perceived/processed as a configural unit, the result can also be explained without appeal to configuration. For example, according to a model such as that of McLaren and Mackintosh [39] the internal activation of X by way of associations with other stimuli present is assumed to produce latent inhibition. When X is presented with B, X can become strongly associated with B and thus receive no internal activation (i.e., no latent inhibition) when without B. However, an X exposed without a unique cue present (e.g., B) would become associated with only the context, assuring the activation of its internal representation when presented, producing latent inhibition. The presence of B during pre-exposure could block (e.g., [27],[28]) or overshadow [54] context-X associations, reducing the internal representation of X evoked by the context and reducing latent inhibition. Such an explanation does not rely on the presence of any unique configural cue, but does depend on the interaction of the stimulus elements by way of within-compound associations between cues.

An explanation based on blocking of context-X associations by B fares less well with the results of Rodriguez and Alonso [61]. Blocking of context-X associations by B should not be affected by the length of exposure to BX. That is, the proportion of X predicted by the context and B together should grow with more exposures, but the relative proportion controlled by each should remain generally constant. Additionally, whether the underlying mechanism is configural or elemental, both frameworks would predict that exposure to a compound such as CX should reduce generalization to AX to a similar extent as exposure to BX, since both compounds include X and could, in principle, support either a configural representation or blocking of context-X associations by C. Yet Sanjuán, Alonso, and Nelson [66] found that preexposure to the compound BX markedly reduced generalization of a conditioned response to AX, whereas pre-exposure to CX had a significantly smaller effect and equivalent to presenting B and X separately. This finding suggests that what is most important is not simply exposure to X in any compound, but exposure to the specific compound that is used in the test. That is, exposure to BX may reduce the perceived similarity between BX and AX, whether through configural differentiation or through changes in associative structure, but the result suggests that preexposure modifies stimulus representations in a stimulus-specific way, and that generalization is sensitive to the history of the exact test stimulus.

### 1.7 Recovery of perceptual effectiveness

An explanation for perceptual learning offered by Hall [17] uses perceptual effectiveness of stimuli to account for the differences observed between alternating and blocked exposure to AX and BX. In accord with Hall’s idea of perceptual effectiveness as, “…the ability of a stimulus to activate its representation” ([17], p. 43) it can be thought of as the ability of an external stimulus to produce an internal representation with the same properties, or the degree to which some internal representation successfully serves as a stimulus. According to his explanation, changes in the perceptual effectiveness of stimuli can occur because of their representations being associatively evoked. For instance, if the salience of a physical stimulus is reduced during exposure (e.g., habituation), Hall suggests that its salience can be restored if the physical stimulus is absent, yet its representation is associatively retrieved by another stimulus. In that case, the effectiveness (in terms of salience in this example) of the stimulus itself would be increased to prior levels upon the subsequent presentation. We will refer to this effect as *restoration of perceptual effectiveness (RPE)*.

In a study by Hall, Blair, and Artigas [18], exposure to BX and X was alternated, followed by exposure to a compound of CX. Here, B and C are equally exposed, but B has had the opportunity for the restoration of its salience on the X trials. Such a restorative effect is not possible with C as it occurred after the X trials. A second group received the same treatment, except that there were no X alone trials when BX was exposed. An aversion to X was subsequently conditioned and the rats were tested with BX and CX. In the condition where there was the opportunity for B to be associatively activated during pre-exposure, there was less expression of the conditioned aversion to X in BX than with CX. In the condition where BX was exposed without the X trials, where B could not be associatively activated in pre-exposure, the aversion to BX and CX was equal. Alternated exposure to BX and X appeared to maintain the salience of B, whereas exposure to only BX did not.

The RPE mechanism, however, cannot operate as effectively after exposure to BX alone, at least not during the preexposure phase. If the RPE mechanism does contribute, it would have to do so during the conditioning phase with AX, where X might associatively retrieve the representation of B. The associatively activated representation of the absent B would restore the salience of the physical B, making its physical presence more effective at affecting test responding. During conditioning of X in the experiments of Hall et al., [18] conditioning of X might enhance the salience of both B and C equally in the conditions where only BX and CX were exposed; there was no control against which to assess that possibility. Nevertheless, the alternated BX/X pre-exposure maintained B’s salience over C.

In the case of exposure to AX and BX, all elements initially suffer a loss in effectiveness. However, when exposure occurs in alternation, the repeated evocation of the representation of A when BX is presented (through its association with X) can lead to a recovery of A’s effectiveness, and the same could occur for B when AX is presented. This recovery process is assumed less likely to occur during blocked presentations of the stimuli where extinction of A↔X should occur during the block of BX presentations (but see; [62], [63]). Thus, during conditioning of AX, stimulus A, who’s effectiveness has been maintained during alternated pre-exposures, is especially salient relative to X and may be better able to overshadow X [36], reducing X’s ability to evoke a response on the test with BX. Also, the presence of a salient B on test, a stimulus not associated with the US, might further produce some decrement in responses controlled by X by way of commanding attention (distraction) or external inhibition [55].

### 1.8 Retrospective revaluation

An additional mechanism to consider in the case of test-stimulus pre-exposure involves the phenomenon of retrospective revaluation (RR). Retrospective revaluation occurs when the associative status of one of two associated stimuli changes due to the acquisition of new information (e.g., conditioning) and that of its associated cue (not present in conditioning) changes in an opposite way.

An example of this phenomenon is seen in backward-blocking designs. For instance, in a human causal judgment task conducted by Chapman [10], participants initially experienced AB+/CD+ training where compounds of stimuli AB and CD were paired with an outcome (+) across several trials. In a subsequent phase, stimulus A alone was paired with the outcome (A+) to increase its associative strength, while stimulus C alone was presented without the outcome (C-) to decrease its associative strength. During the final test, stimuli B and D were presented. Stimuli B and D were not present during the second phase and no direct changes in their associative strength were expected. Nevertheless, B was rated as less effective than D in predicting the outcome. It is as if the associates of A (B) and C (D) were retrospectively re-valued in the second phase.

In the experiments that show perceptual learning about one stimulus, retrospective revaluation might occur. For example, during pre-exposure to BX, B and X may become associated. During subsequent conditioning of AX, the representation of B would be retrieved by X. If the retrieval of B changes its associative properties in a manner opposite that of X’s association, then B, which began as neutral, may become inhibitory for the outcome (e.g. [74]). Thus, the response to BX on test would be reduced by way of B having acquired some outcome inhibition through a retrospective-revaluation process.

Under similar conditions, however, retrospective revaluation is not always observed, but rather mediated conditioning effects are sometimes obtained. For example, in an experiment by Holland [23], a tone was initially paired with food, establishing an excitatory association. The tone alone was then presented and followed by lithium chloride, an aversive stimulus. In theory, the tone should evoke a representation of food at this time, which was being paired with the lithium chloride. During the test, animals consumed less food when it was preceded by the tone, especially after the aversive pairing. Shevill and Hall [70] showed that following exposures to a compound stimulus AX, extinction of A following conditioning of X led to a slight reduction in responding to X (mediated extinction).

Many similar examples exist in the sensory pre-conditioning literature (e.g., [7], [60],[78]). There, two neutral stimuli are presented without an outcome (e.g., AB-) and then one of those is subsequently reinforced (B+), which results in a response to A being observed. Sensory pre-conditioning might occur by way of mediated conditioning where A is associatively evoked by B and, thus, paired with the outcome, or by way of an associative chain where A evokes B which evokes the outcome expectation. In experiments where the contribution of associative chains between B and A are minimized while those between A and B are maintained (e.g., A predicts B, but not vice-versa), retrospective revaluation is observed when A’s associative status is modified [70].

Research investigating the critical manipulations separating when one phenomenon occurs as opposed to the other has failed to provide clear characterizations. Nevertheless, both occur. Although there are studies that show revaluation in animals (e.g. [29], [30], [40]) this phenomenon is particularly evident in human studies (e.g., [1],[10],[11],[33],[37],[68],[80]vogel 75), using simultaneous compounds (see [10]; Larkin et al, 19998[36]).

Theories, such as Wagner’s SOP model [76] explain associative changes in stimuli based on their states of activation. According to Wagner, stimuli pass through three states of activation: A1, A2, and I. The A1 state is produced by the environmental presence of the stimulus itself and represents a highly active level of activation where a stimulus might be vividly experienced. This state changes over time to an A2 state, which can persist after the stimulus is no longer present before decaying to a state of inactivity (I), where the stimulus no longer should influence perception or behavior. Once a stimulus decays to A2, it cannot be re-activated into the A1 state until it has passed to inactive.

Wagner’s model posits that when stimuli are in the primary A1 state simultaneously, excitatory associations form between them. When two stimuli are associated in this manner, the predicting stimulus in A1 can activate the predicted stimulus into its A2 state. It is from this state that behavior associated with the expected stimulus occurs. However, when a CS is in the A1 state and an outcome is in the secondary A2 state (e.g., it is predicted, but does not occur, such as in extinction), an inhibitory association between the CS and the outcome is established. When the CS, or both stimuli, are in A2, no learning occurs. Thus, an element such as “B” that is associatively activated to A2 by way of within-compound associations is less able to be learned about. This limiting of learning when stimuli are predicted by way of within-compound interactions is a property incorporated, in different ways, in many subsequent theories (e.g., [21], [22], [39], [69]).

To address retrospective revaluation, Dickinson and Burke [11] proposed a simple modification of the SOP model that, unlike Wagner’s original framework, allows associative changes to occur for stimuli that are absent but retrieved associatively (i.e., in A2). According to their proposal, inhibition forms between two stimuli when they are in different states of activation. While Wagner’s model suggests that no learning occurs when a CS is in A2 and an outcome is in A1, Dickinson and Burke argue that just as an inhibitory CS-Outcome association can occur when the CS is in A1 and the outcome is in A2 (as in extinction), an inhibitory CS-Outcome association can also form under when the CS is in A2 (expected, but absent) and the Outcome in A1. This modification implies that the associative retrieval of a stimulus, placing the representation of an absent element into the A2 state, can facilitate the formation of an inhibitory link between that stimulus and a present outcome.

The modified SOP model by Dickinson and Burke [11] can be easily applied to explain the perceptual learning effects discussed here. During exposure to BX, strong within-compound associations between B and X are likely established. When AX is subsequently conditioned, B can be retrieved into A2 through the shared element X. B begins with no association with the outcome, thus the mismatch in states between B (A2) and the outcome (A1) lead to B acquiring inhibition. Consequently, during the subsequent test with BX, an inhibitory B would diminish the expression of any associative strength acquired by X.

The present experiments were conducted to assess the contributions of retrospective revaluation and RPE to the enhanced discriminability of the test-stimulus BX following exposure. If exposure to BX allows the evocation of B during conditioning with AX, B might become inhibitory as Dickinson and Burke [11] predict, or B might recover its otherwise lost salience increasing the discriminability of compounds containing it [17]. Following Rescorla [59], the potential properties of B were examined with summation (Experiment 1a) and retardation (Experiment 1b) tests.

## 2 Experiments 1a & 1b

Participants were volunteers from Prolific Academic. As anonymous members of research-participant platform [58] they are aware that they are participating in research when they select a task for which Prolific has informed them that they qualify. The study description informed potential volunteers of the requirements of the research project (a computer capable of conducting the task), that their participation was voluntary, and that they could withdraw their data from the project after completion. They were explicitly informed that by clicking the link which directs them to the experiment they were providing their consent to participate. There are no minors participating in the Prolific platform.

The experiments (Exp 1a & 1b) were conducted online using the cited platform. Both used a video game, developed based on an earlier version [46], which has been shown to be an effective tool for investigating perceptual learning ([47], [48], [67]) and associative learning phenomena in general (e.g., [51],[45],[52]). All procedures were reviewed and approved by the Ethics Committee for Research on Human Subjects (project, 2024/058, reference number M10/2024/079 Rev2) and conducted in accordance with the ethical standards of the Helsinki Declaration and its later amendments.

In the method, participants earn points by shooting torpedoes at a “cybergnostic space chicken” to pass a “promotion” exam. During the game, they can be attacked by the chicken with an “egg of destruction” which causes them to lose points. They are instructed that they can avoid point loss by suppressing their own rate of torpedo launching to save power immediately prior to the attack. Colored sensors can be programmed to appear which signal the attack in the conditioning phases, and participants learn to suppress their mouse clicking (torpedo firing) during the presence of the sensor in anticipation of the attack to avoid point loss.

The design is shown in Table 1. The study employed a completely within-subjects design in which participants underwent two phases of training and a test. The first phase involved exposure to compound stimuli. Each oval sensor was composed of two colors (e.g., BX), with the color represented by the first letter in a compound’s name (e.g., B) occupying 16% of the sensor, from left to right, while the color represented by the second letter (e.g., X) occupying the remaining 84%. Compounds BX, CY, and DZ were exposed 6 times each. The second phase focused on conditioning with two additional stimulus compounds (AX & AY) each containing elements (X & Y) which had been associated with elements from pre-exposure. AX and AY were followed by the egg-of-destruction attack (+), representing the unconditioned stimulus (US).

**Table 1.**
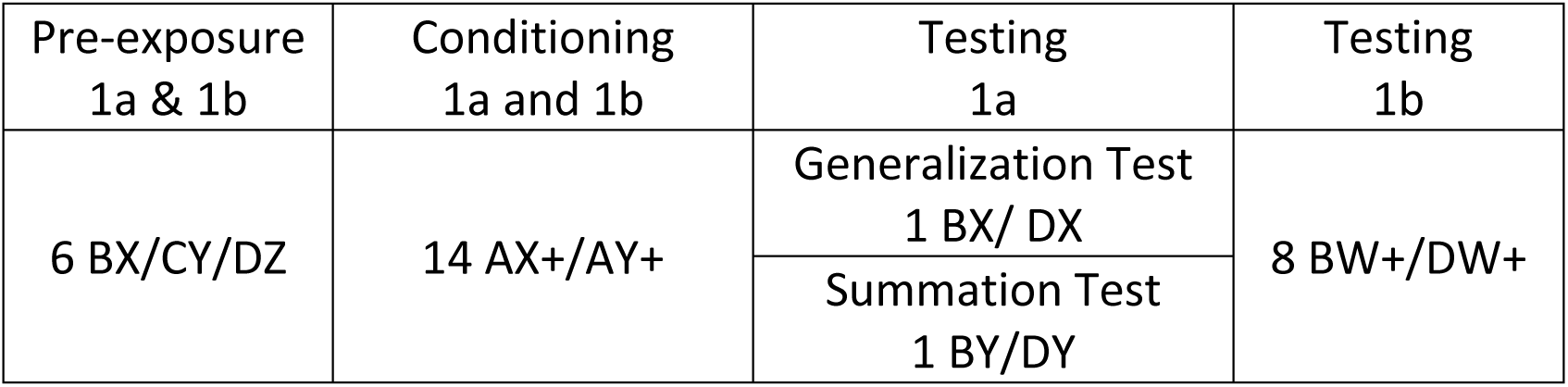
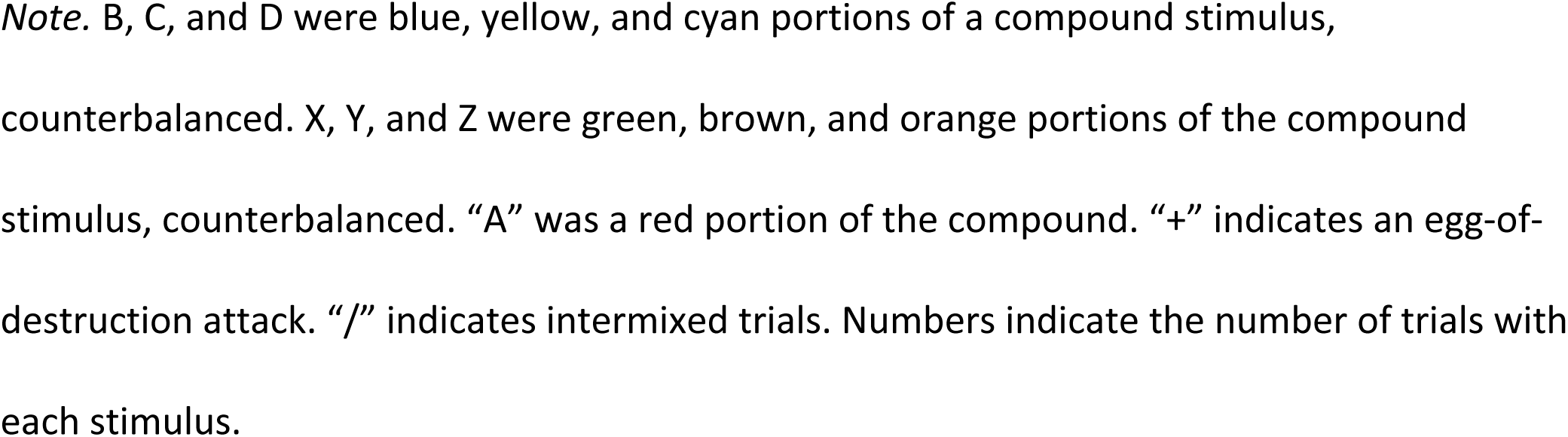
Design of Experiments 1a and 1b.

| Pre-exposure<br>1a & 1b | Conditioning<br>1a and 1b | Testing<br>1a | Testing<br>1b |
| --- | --- | --- | --- |
| 6 BX/CY/DZ | 14 AX+/AY+ | Generalization Test<br>1 BX/ DX | 8 BW+/DW+ |
|  |  | Summation Test<br>1 BY/DY |  |

*Note.* B, C, and D were blue, yellow, and cyan portions of a compound stimulus, counterbalanced. X, Y, and Z were green, brown, and orange portions of the compound stimulus, counterbalanced. “A” was a red portion of the compound. “+” indicates an egg-of-destruction attack. “/” indicates intermixed trials. Numbers indicate the number of trials with each stimulus.

During pre-exposure, B and X can become associated. Thus, during conditioning with AX there is an opportunity for X to associatively activate B, making it inhibitory through RR, or restoring its perceptual effectiveness. Stimulus D, also present in a compound during pre-exposure, has no opportunity to be associatively activated as its associate, Z, is never presented again. Thus, D cannot be affected by either RR or RPE. The effects of conditioning B’s associate were examined using the summation and retardation tests outlined by Rescorla [59] in Experiment 1a and 1b, respectively.

### 2.1 Experiment 1a

In Experiment 1a training was followed with compound tests. Generalization with BX and DX was assessed along with the inhibitory properties of B by comparing BY and DY. The tests each compare two stimuli, B and D, both of which have been equally exposed in a compound but vary only by whether they had the opportunity to be associatively activated prior to testing.

The associative activation of B could allow for retrospective re-valuation to occur making B inhibitory. According to Hall [17] the effectiveness of stimulus B, which is otherwise being habituated, would be restored. Based on previous findings with a similar method ([47],[48],[50],[67]), we anticipated a perceptual-learning effect where the conditioning of AX would generalize less to BX than to DX, despite equal exposure to both B and D in compounds. If conditioning with AX allows the activated memory of B, through X, to become inhibitory for the outcome through RR, suppression to BY (inhibitor + excitor) should be weaker than the response to DY (neutral + excitor). The same outcome on the BY/DY summation test would be expected based on Hall [17]. The restored effectiveness of B would make it functionally more salient and distracting on test relative to D, better reducing suppression controlled by X by way of external inhibition [55]. If some configural process or unitization is responsible for the reduction in generalization to the test stimulus, then we would expect a larger effect with BX vs DX than BY vs DY, as BX is the original exposed compound.

### 2.2 Experiment 1b

Experiment 1b was conducted using the same method and design in the pre-exposure and conditioning phases. Rather than summation and generalization tests, Exp 1b concluded with a retardation test where B and D were combined with W (white), on separate trials, and paired with an egg attack. The summation and retardation tests are seen as necessary tests for detecting conditioned inhibition [59], and both are particularly relevant in this experimental design. During the summation test, B could inhibit suppression evoked by Y because it is inhibitory for the outcome (conditioned inhibition) by way of retrospective revaluation. In the case of B being inhibitory, acquisition of suppression in the retardation test should be *less* rapid in compounds involving B. Or, as mentioned earlier, B could affect X and Y by way of external inhibition. B could be especially salient and more distracting than D if B was associatively activated during AX conditioning and its effectiveness restored, as predicted by Hall [17]. In that case, a compound of BW should be more salient than DW and acquire suppression *more* rapidly.

## 3 Method

### 3.1 Participants

All procedures involving human participants were reviewed and approved by El Comité de Ética para las Investigaciones relacionadas con Seres Humanos (project, 2024/058 Extinción, inhibición y cambios representacionales en el control contextual, reference number M10/2024/079 Rev2) and conducted in accordance with the ethical standards of the Helsinki Declaration and its later amendments.

The apparatus used here was also used by Nevado and Nelson [52] to investigate the “renewal effect” (e.g., [51]). Extinguished conditioned responding increases with a change in the background context where extinction occurred. In that online within-s experiment Nevado and Nelson obtained *η^2^_p_* = .25 for the renewal effect, which is our current best estimate as to the maximum effect we might expect to observe. Counterbalancing the different identities of the stimuli required 36 different combinations. The method updates a log when each participant is assigned, keeping conditions balanced. However, participants may get assigned but not complete the task, resulting in the need to recruit again. We planned to recruiting 100 participants to ensure that we would fill all 36 conditions at least twice through random assignment, and have adequate power. Power was .17, .70, and .98 to detect Cohen’s [9] small, medium, and large effects, and .8 to detect *η^2^_p_* = .075, (one-third of the estimated maximum effect). The effect observed in Experiment 1a produced *η^2^_p_* = .05. For Experiment 1b N was increased to plan on recruiting 150 participants to have a power of .8 to detect that effect size.

Participants were randomly assigned to counterbalancing conditions without replacement until all conditions were filled and then re-entered into the pool. Participants might begin the experiment and be assigned, updating the pool, but not complete the study. For that reason, more slots than needed were available resulting in the recruitment of 102 people in Exp 1a and 165 in Exp 1b. The age and gender statistics of the sample, based on those who reported such demographics through Prolific, are shown in Table 2. The distribution of gender was independent of the experiment, Χ^2^ = .34, *p* = .56.

**Table 2.** Sample demographics.

| Experiment | Reported Gender | N Reporting | Min. age | Max age | Mean | Sd |
| --- | --- | --- | --- | --- | --- | --- |
| 1A | Male | 62 | 20 | 80 | 41.29 | 14.13 |
|  | Female | 40 | 19 | 68 | 37.05 | 13.41 |
| 1B | Male | 103 | 18 | 73 | 37.61 | 12.48 |
|  | Female | 57 | 19 | 66 | 39.92 | 12.60 |

*Note.* Table shows the distribution of self-reported gender by experiment. It also reports the mean, standard deviation, minimum and maximum values for age.

### 3.2 Apparatus

A detailed description of the video game used in the experiment can be found in Nevado & Nelson ([52]; see also [49]). The apparatus summarized here was the same as what is described in more detail in Nelson and Sanjuan. Participants played a video game where they were cadets taking a test to earn a promotion. To pass the test they had to earn points by shooting a “cybergnostic space chicken.”

The apparatus is shown in Fig 1. Participants interacted with a video game that simulated the experience of flying in a spaceship, offering a first-person view through a view screen into a colorful 3-d galaxy. The interface featured a dark gray gun attached to a pillar at the top of the screen. The gun tracked a red circular crosshair which was moved by the participant’s mouse. Key information, such as the current context’s name (Boutonia, which was not manipulated in the current experiments), was displayed in a translucent label on the top left of the screen. Another panel was located at the top right of the screen that displayed either “Gaining Points,” “Not gaining points,” or “points lost” (with the number of points lost displayed in the blank). Instructions (detailed in [52]) were presented on a translucent black panel that rose from the bottom of the screen.

**Fig 1.**
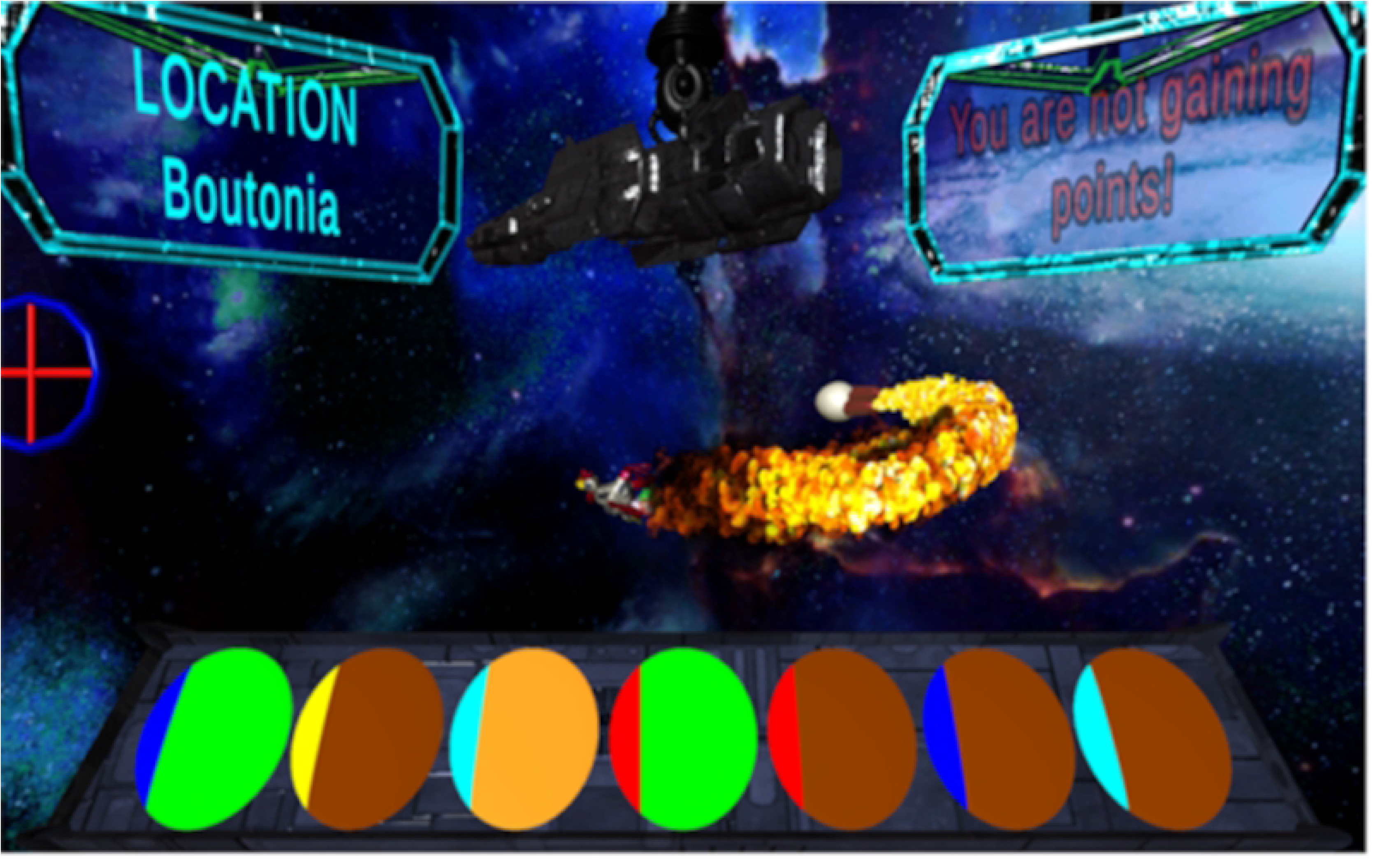
Screenshot of the apparatus. Figure 1 shows the apparatus and example stimuli, from left to right, BX, CY, DZ, AX, AY, BY, DY. Blue, Yellow and Cyan were counterbalanced as B, C, and D. Orange, Green, and Brown were counterbalanced as X, Y, and D (36 conditions). Though shown in different positions here, all CSs were presented in the middle position (fourth from the left). In the image, the Chicken has launched an egg of destruction (the US).

**Fig 2.**
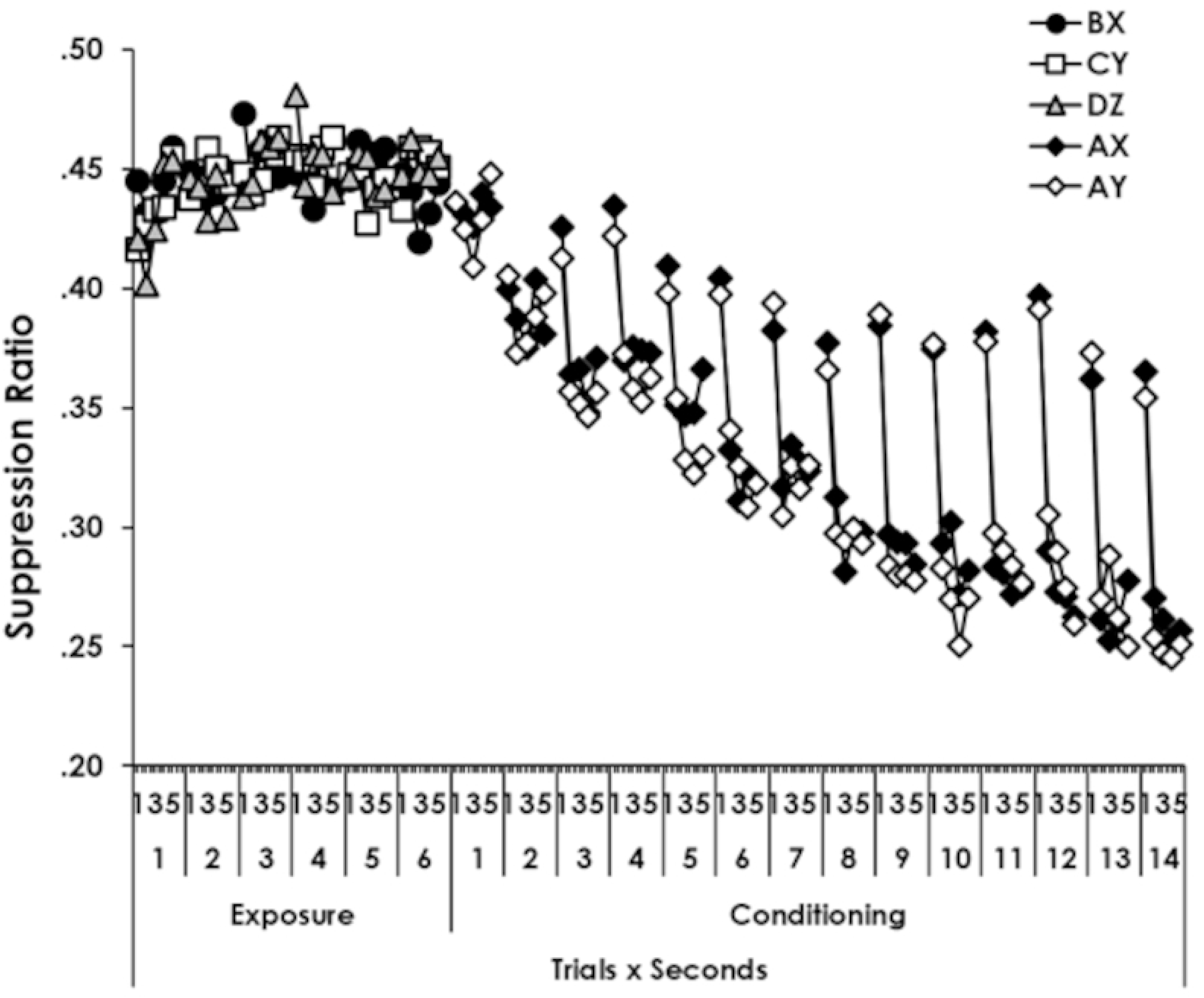
Suppression during pre-exposure and conditioning. Figure 2 shows suppression to the stimuli on each second of each trial during exposure and conditioning for Experiments 1a and 1b combined. Error bars are omitted to remove clutter. Notice that the Y axis begins at .2.

Participants began by entering their Prolific ID and pressing “B” to start the game, which commenced in a “training context” that appeared as a green wireframe cube. Initial instructions were delivered both as on-screen text and through a pre-recorded male voice. Participants were guided through a brief tutorial that explained how to operate the gun, charge the weapon by clicking the mouse at a rate of three clicks per second, and maintain a steady rate of firing at the chicken.

The tutorial emphasized the importance of interpreting in-game events, such as flashing lights or sounds, to predict “egg of destruction” attacks by the chicken. Participants were instructed to reduce their firing rate when anticipating an attack to conserve power and minimize point loss. Following the instructions, participants completed a short multiple-choice quiz designed to ensure their understanding. Questions were repeated until answered correctly, with incorrect responses prompting a “wrong answer” message in red and a repeat of the question. Correct answers were followed by a feedback screen indicating the answer was correct and reiterating why the answer was correct.

After the tutorial a robotic chicken appeared together with a music background. This character was animated with visual and audio features, including randomly varying movement patterns and robotic chicken sounds. The chicken’s behavior was governed by several randomly changing variables, such as speed, direction, and duration of movement, ensuring unpredictable interactions based on the chicken’s behavior. The view of the galaxy changed as the camera tracked the movement of the chicken in 3d space.

During the game, eight different stimuli were used, consisting of various color combinations flashing (fading on and off, cycling at 3 Hz) on the center oval of the seven black ovals appearing in the trapezoidal panel the bottom of the screen. The unconditioned stimulus (US) was an “Egg of Destruction.” This egg was visibly launched from the rear of a chicken, accompanied by a burst of flame from the rear of the chicken. The egg, propelled by visible flames from two attached exhausts, traveled rapidly until it struck the participant’s screen, resulting in a large fireball explosion accompanied by a deep explosion sound and electrical after effects. The time from the egg’s launch to its impact varied depending on the chicken’s distance from the screen, but the entire sequence, from launch to explosion, was completed in no more than 4 seconds.

A “test screen” could be presented which was a screen of static presented for 5-s with the words ““Test control has no information on the outcome at this time. Simulation will resume shortly.” The text appeared in the center of the screen in a blue font. This screen was used as the outcome on test trials to help avoid carry-over effects in testing [32]. The screen also appeared in training, as indicated in the procedure below, to familiarize the participants with its occurrence.

Sensor colors were composed of a left-most (left 16% of the sensor area) and right-most (right 84% of the sensor area) color. The colors appearing in the left-most of the sensor were B, C, D, and A while X, Y, and Z colors were in the right-most part. “A” was always Red while B, C, and D were Blue, Yellow, and Cyan, counterbalanced. Colors X, Y, and Z were Green, Brown, and Orange, counterbalanced fully against B, C, and D. When presented, the sensors always appeared in the fourth from the left of the seven ovals. Examples are shown in Fig 1, though all stimuli appeared in the center oval in the experiments.

### 3.3 Procedure

#### 3.3.1 Experiments 1b & 1b

Participants engaged in three experimental phases: Exposure, Conditioning, and Testing. During the Exposure phase, BX, CY, and DZ were presented one time each in each of 6 blocks. Each presentation lasted 5 seconds and the order of presentation within a block was randomized. There was no outcome following the presentations, except on block 3 the test-screen outcome appeared following each stimulus. The test-screen outcome was included in each phase to familiarize participants with its occurrence.

Participants were firing at the chicken during this and all subsequent phases. No other specific events occurred in phase 1. The inter-trial interval (ITI) was randomly determined on each trial in each phase from a flat distribution with a mean of 11 seconds and a standard deviation of S = 5 and clipped to a range of 5-17 (resulting ***x̅*** = 10.98, S = 4.57).

The conditioning phase began uninterrupted. Colored sensors AX and AY each appeared 14 times and were each paired with an “Egg of Destruction” attack. There were 7 blocks of four trials containing two AX and two AY trials in each block, with the trial order randomly determined. The attack drained up to 100 points based on “<u>d</u>amage” which was a positively accelerating function of their suppression (see [52], for a full description of the mapping of suppression to damage). No suppression would result in maximum damage (d = 1) and a loss of 100 points (i.e., d × 100), while complete suppression would result in no damage (d = 0) and no loss of points. One trial with both AX and AY was followed by the test-screen outcome on both the 2^nd^ and 6^th^ blocks to ensure that the test screen was not associated with any particular phase.

#### 3.3.2 Experiment 1a: Compound Testing

Compound tests were used to assess generalization and inhibition. Generalization from AX was assessed with a single presentation of BX and one of DX, each followed by the test-screen outcome. The order of the two trials was randomized. The inhibition assessment was a summation test consisting of a single presentation of BY and one of DY. The order of the two trials was randomized. The order of the summation and generalization tests was randomized.

#### 3.3.3 Experiment 1b Retardation testing

Experiment 1b followed conditioning with a retardation test where B and D were combined with W (white occupying the right-most portion of the sensor). We reasoned that simply using B and D, with the remaining portion of the sensor black, may promote attention to these stimuli, obscuring any differences that the pre-exposure manipulation may have produced. There were 16 trials arranged in four blocks of four trials. Each block contained 2 BW+ and 2 DW+ trials with the order of the four trials randomly assigned.

### 3.4 Data analysis

The number of mouse clicks during the ITI and CS were converted to rates (clicks per second) and used to calculate standard suppression ratios as CS rate / (ITI rate + CS rate). Suppression was analyzed using repeated measures Analysis of Variance (ANOVA) and Type III sums of squares by IBM SPSS v23 (SPSS, 2015). Effect sizes are reported as partial eta squared (*η^2^_p_*) and 95% confidence intervals were calculated using non-central F distribution functions for Excel found in Nelson [42].

Participants suppress very little during the first second of the CS presentation in this method online, and suppression tends to asymptote around .2 (see [52], [49] c.f., [51]). To increase the sensitivity of the measure, we analyzed suppression on a per-second basis of each second of the CS. In the method upon which the present method was conceptually modeled, suppression during the CS has been shown to increase over the duration of the CS [43], revealing effects that otherwise may go undetected.

The first two phases were analyzed with 1a and 1b combined, and “Experiment” (1a vs 1b) included in the analysis. The test data were analyzed by experiment. Suppression can only be calculated when ITI rates of responding are above zero. The occasional zero ITI rate was replaced with the mean of the participant’s other non-zero trials of the phase. We followed previous procedure used with this method [52] and removed participants from the analysis of a phase when 20% or more trials were replaced in that phase.

### 3.5 Transparency

The data are available at in .sav format for SPSS and .xlsx format for Excel. Analysis syntax is available in text files with .sps extension for use in SPSS.

## 4 Results

There were no differences between Experiments 1a and 1b in the training data, which are combined and shown in Fig 1. The figure shows suppression on each second of the presentation of each CS. Data from pre-exposure are shown at left above “Exposure” and conditioning of AX and AY are shown above “Conditioning.”

### 4.1 Exposure

Data from exposure were analyzed with a CS (BX, CY, DZ) x Trials x Seconds x Experiment ANOVA. No participant was removed by the ITI-rate screening. The analysis showed a Trials x Seconds interaction, *F*(20, 5280) = 1.6, *p* = 0.04, *MSE* = 0.03, *η*^2^*_p_* = 0.006, *CI*_95%_ = 1.9 × 10^-8^ - 0.007. Inspection of the figure suggests that suppression decreased during the seconds of the first trial, but randomly varied among other trials. Of most importance, there were no effects involving CS or Experiment. The Trials x Seconds effect reported above was the only reliable effect in the analysis, *p*s ≥ .11.

### 4.2 Conditioning

Data from conditioning were analyzed with a CS (AX vs. AY) x Trials x Seconds x Experiment ANOVA. Six participants were removed by the ITI screening. The analysis showed effects of Trials and Seconds which were superseded by a Trials x Seconds interaction, *F*(52, 13416) = 4.81, *p* = 2.4 × 10^-27^, *MSE* = 0.03, *η*^2^*_p_* = 0.02, *CI*_95%_ = 0.011 - 0.02. The pattern is clear in the figure. Suppression was acquired over trials, and increased across seconds within each trial with that increase being greater in later trials than in earlier ones. There were no other effects, *p*s ≥ .17.

### 4.3 Compound Testing (1a)

The data from the Compound Testing are shown in Fig 3. Compounds assessing generalization (BX and DX) are show at left and the those assessing inhibition (BY and DY) are shown at right. Suppression on each second of the CS presentation is presented.

**Fig 3.**
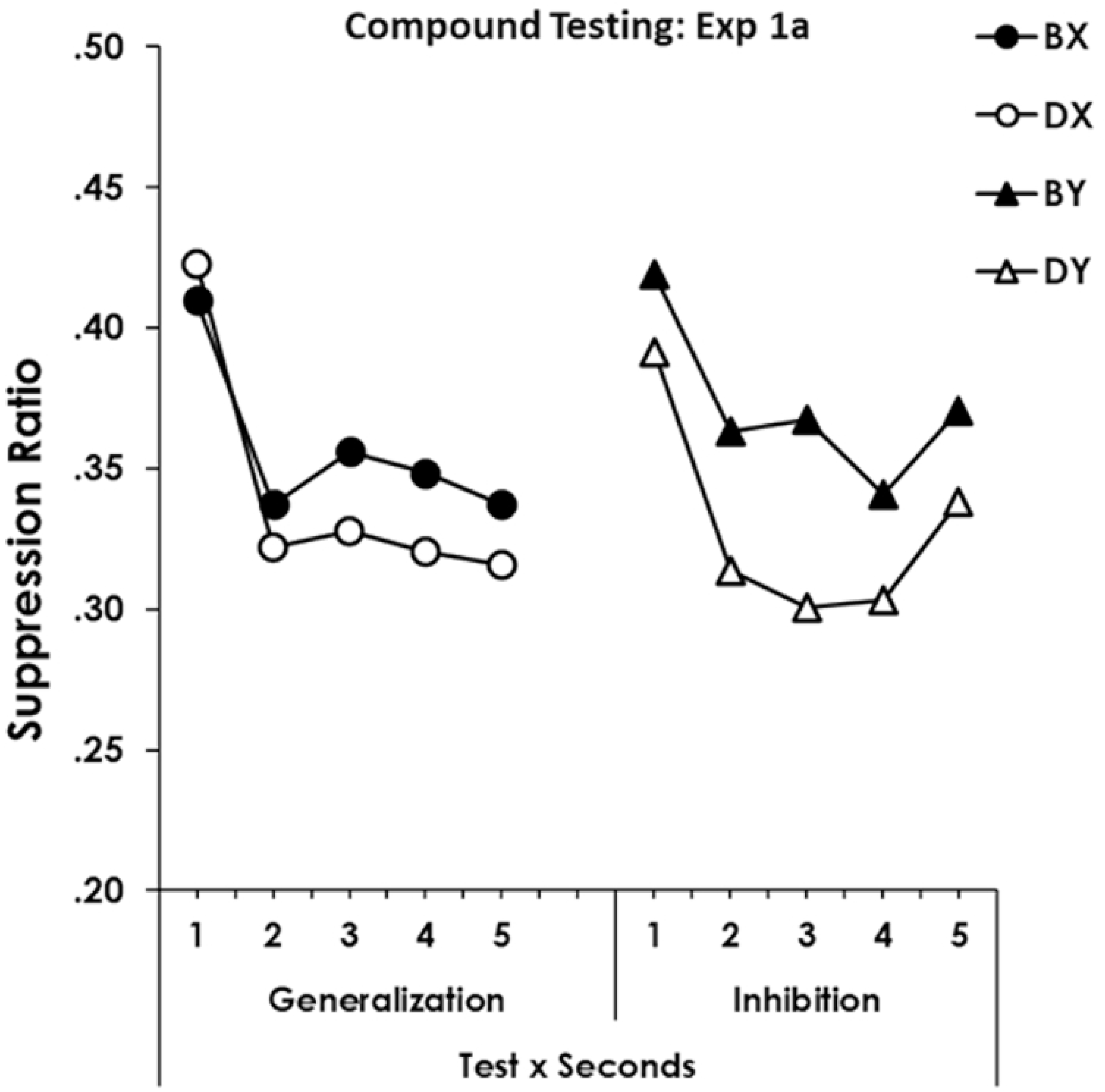
Generalization and summation testing. Figure 3 shows suppression to BX and DX in the generalization test at left and BY and DY from the inhibition testing at right. Error bars are omitted to remove clutter. Notice that the Y axis begins at .2.

The data were analyzed with a Compound-type (Generalization involving X vs. Summation involving Y) x Revaluation (B vs D) x Seconds ANOVA. Five individuals were eliminated by the ITI screening. A retrospective-revaluation effect would appear as a reduction in suppression to BX relative to DX and a parallel reduction in suppression to BY relative to DY. A lack of a retrospective-revaluation effect would produce either no effects, or an interaction of Compound-Type with Revaluation. The analysis produced a Revaluation main effect, *F*(1, 96) = 5.07, *p* = 0.03, *MSE* = 0.08, *η*^2^*_p_* = 0.05, *CI*_95%_ = 1.0 × 10^-6^ - 0.16. There was no effect of Compound-Type, *F* < 1, and no Compound-Type x Revaluation interaction, *F*(1,96) = 1.24, *p* = .27. There was an effect of seconds, *F*(4, 384) = 11.97, *p* = 3.5 × 10^-9^, *MSE* = 0.04, *η*^2^*_p_* = 0.11, *CI*_95%_ = 0.05 - 0.17, but no interactions of seconds with the other variables, *p*s ≥ .41. Conditioning the exposed CS (B), whose associate had been conditioned, led to a greater disruption of responding in a compound than did D that had been exposed and whose associate was not experienced again.

### 4.4 Retardation Testing 1b

The retardation test data are presented in Fig 4 which shows suppression on each second of each trial to the BW and DW stimulus compounds. The right-most portion shows the effects of Trials and Compound type, collapsed over seconds.

**Fig 4.**
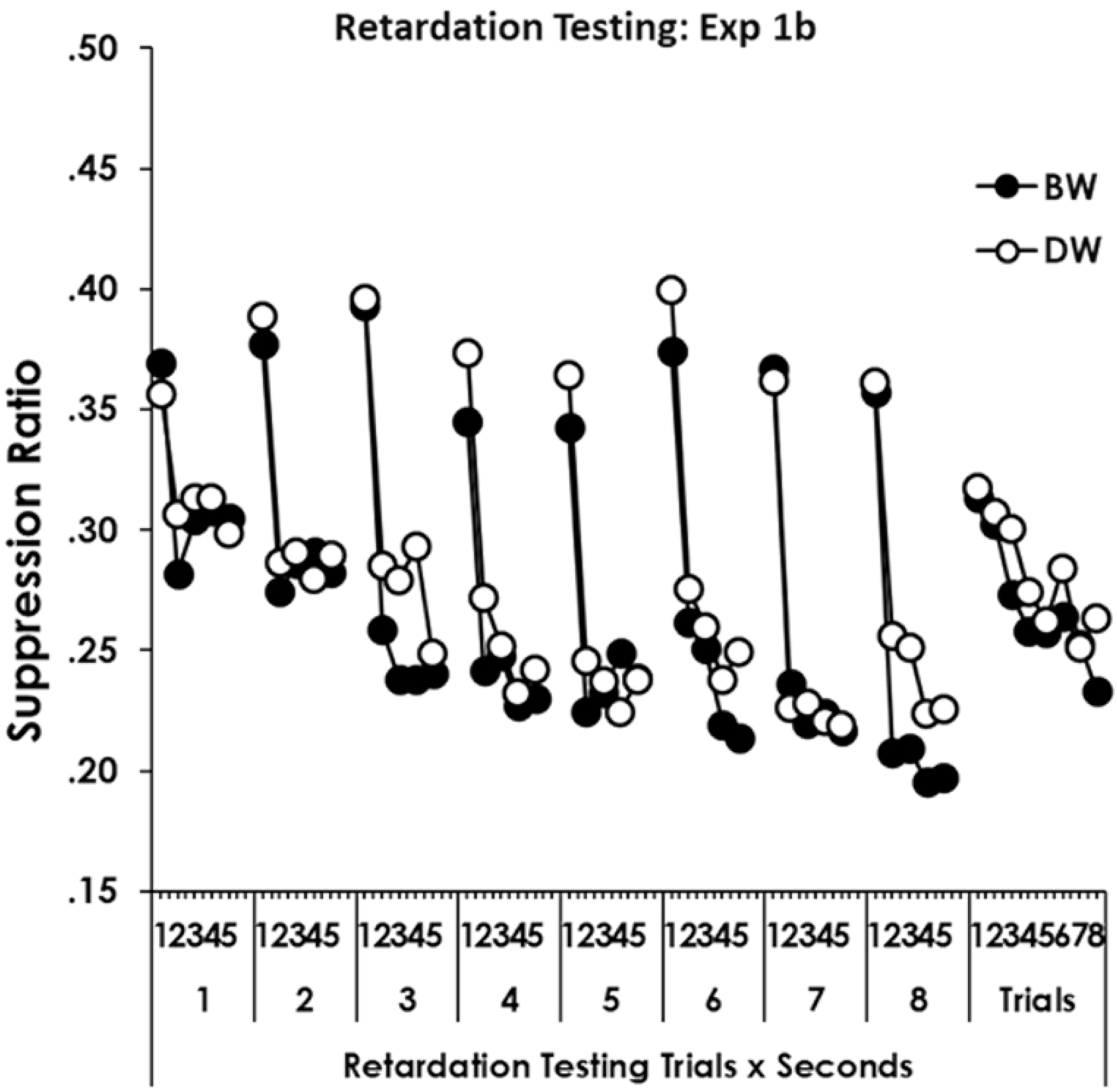
Retardation testing experiment 1b. Figure 4 shows suppression (Y axis begins at .15) to BW and DW on each second of each trial above *Retardation Testing Trials x Seconds*, and shows suppression averaged over seconds on each trial above *Trials*. Error bars are omitted to prevent clutter.

The data from retardation testing were analyzed with a Compound Type (BW vs. DW) x Trials x Seconds ANOVA. Eight persons were excluded by the ITI response screening. The analysis showed a main effect of Compound Type where suppression to BW was slightly greater than DW, *F*(1, 150) = 8.12, *p* = 0.005, *MSE* = 0.06, *η*^2^*_p_* = 0.05, *CI*_95%_ = 0.005 - 0.13. There were effects of Trials and Seconds which were superseded by a Trials x Seconds interaction, *F*(28, 4200) = 2.8, *p* = 1.3 × 10^-6^, *MSE* = 0.02, *η*^2^*_p_* = 0.02, *CI*_95%_ = 0.006 - 0.02. The Trials x Seconds effect is clear in the figure where suppression increased across seconds of the CS presentation, with that effect being more pronounced later in training. There were no further effects, *p*s ≥ .64.

## 5. General discussion

How does experience with one stimulus enhance its discriminability from another, similar stimuli encountered later? The primary goal of this study was to better understand the mechanisms underlying this process of perceptual differentiation, with a particular focus on evaluating two potential explanatory mechanisms. The study investigated whether retrospective revaluation (e.g., [11]) or a recovery of perceptual effectiveness (e.g. [17],[18]) can contribute to the enhanced discriminability observed when a test stimulus has been pre-exposed. In both experiments experimental (B) and control (D) elements were pre-exposed in compounds (BX and DZ). Following pre-exposure, compounds AX and AY were conditioned. Conditioning with AX allows for the associative activation of B which may make it inhibitory by way of RR, or have its otherwise habituated effectiveness restored (RPE). Neither of these processes can operate on D for which its associate Z is never encountered again.

In Experiment 1a the conditioned X element from the AX compound was tested in compound with B, matching the pre-exposed BX, or it was tested with D, an equally pre-exposed stimulus, but one which produced a novel compound. B and D were also tested with Y where neither compound matched pre-exposure. If B were inhibitory through RR, or more perceptually salient through RPE, there should be less suppression to BX than DX, as well as less suppression to BY than DY. If the generalization-reducing effect depended on some BX configural cue produced in pre-exposure, then an effect would only be observed with BX vs. DX, but not BY vs. DY. The results were clear; the effects were dependent on B. There was an effect of whether an element of the test stimulus could have been associatively activated earlier, producing either RR or RPE, that applied equally to tests involving the pre-exposed test stimulus or those involving new compounds with the pre-exposed element as indicated by the lack of an interaction [14],[53].

The retardation test of Experiment 1b determined whether the mechanism of action observed in 1a was RR or RPE. An RR interpretation would assume B was inhibitory for the outcome, thus a compound of BW should condition more poorly than DW. A RPE interpretation would assume the more effective representation of B might commanded more attention than D, thus a compound of BW would condition better that DW. The results were again clear. Conditioning with BW was more effective than with DW, supporting the idea that B was effectively more salient than D, producing a more salient BW compound than DW. The results support RPE as they are consistent with the idea that B was more salient than D, allowing it to pass a summation (Experiment 1a) test, while failing a retardation (Experiment 1b) tests for inhibition.

Importantly, a recovery of perceptual effectiveness through associative activation differs from retrospective revaluation in that it places the effect in perceptual terms, rather than in terms of the associative status of a stimulus. The activation of B by X during AX conditioning might restore a representation of B that had diminished in salience during pre-exposure. The perceptual system, in this case, could treat the associative activation of B as a signal to “refresh” or recover the stimulus’ prominence, making it more effective in subsequent presentations, such as in the summation test. This restoration of perceptual salience may be an attentional process whereby the absence of otherwise represented stimuli redirects attention to those stimuli, rather than a process of associative learning that modifies the stimulus’s connection to an outcome (either through excitation or inhibition). Salience recovery may be a consequence of associations, but serves as a non-associative adjustment that enhances the influence of the stimulus in future encounters, while leaving its associative connections unchanged.

Building on this analysis, some differentiation between similar stimuli would not require that one of the stimuli undergoes conditioning to acquire meaning, as required by an RR interpretation. Instead, simply encountering a stimulus may be sufficient to associatively activate the distinctive elements of similar stimuli and restore their perceptual salience. These perceptually effective unique elements enable the organism to discriminate between the stimuli. In this respect, our interpretation aligns completely with Hall’s [17] proposal and evidence showing that the shared element must associatively activate the representation of the unique element to recover its salience. The proposal expands Hall’s analysis in that Hall’s discussion of this process focuses exclusively on preexposure and research exploring this idea have evaluated conditions where activation of the unique element of a stimulus is produced during pre exposure. Our research here shows that this salience recovery can occur not only during preexposure, but at any moment when there is an opportunity to reactivate the unique features of the stimuli.

The presence of B, which conditions would permit its associative activation during conditioning of AX, reduced generalization from AX to BX more than that produced by an equally exposed cue, D, whose associate was not conditioned. B also reduced generalization from AY to BY better than D, and a compound of BW conditioned better than did a compound of DW. These findings suggest that B was more effectively salient than D, producing more distraction or “external inhibition” on the BX test, and was easier to learn about during the retardation test.

These findings contribute to the broader literature on perceptual learning and associative learning. While previous studies have emphasized the role of alternating exposure to stimuli in promoting perceptual learning, the present results demonstrate that exposure to a single stimulus compound (BX) is sufficient to reduce generalization in well-controlled within-subject designs. Differential latent inhibition to common and unique elements, along with mutual inhibition between unique elements that can form with exposure to both training and test stimuli undoubtedly contribute to perceptual learning effects. In the present research, those were either held constant (latent inhibition) or were unable to form (mutual inhibition), thus the effect observed here cannot be explained by those mechanisms. The absence of those mechanisms may account for the small effect we observed, further reinforcing that restored salience is a contributing factor to perceptual learning, but certainly not the only factor.

How do we recognize that a new coffee is different from one previously experienced? A coffee has many flavors within it. Some of those flavors, such as a degree of bitterness, are common to all coffees while other flavors, such as nutty or chocolate notes, are unique to different coffees. Experience with a coffee establishes associations between those elements. When a new coffee is experienced, the common element retrieves a representation of the unique element from the first coffee, bringing the distinctive features of the first coffee back into focus. It would be at this moment that our perceptual system achieves differentiation between the two coffees. This process might restore the perceptual effectiveness of those unique features, making discrimination even simpler with later encounters. The distinctive characteristics of the first coffee might become more salient, allowing them to be better compared with other coffee attributes.

Like the work of Hall et al., [18], these findings suggest that exposure effects involve complex associative mechanisms that affect perceptual processes. By uncovering the roles of both associative retrieval and salience recovery, this work contributes to a more comprehensive understanding of the interplay between perceptual and associative learning.

